# Altered epithelial development of the lateral ventricle choroid plexus in *Mllt11* mutants

**DOI:** 10.1101/2025.02.27.640680

**Authors:** Sam Moore, Danielle Stanton-Turcotte, Emily A. Witt, Angelo Iulianella

**Affiliations:** Department of Medical Neuroscience, Faculty of Medicine, Dalhousie University, and Brain Repair Centre, Life Science Research Institute 1348 Summer Street, Halifax, Nova Scotia, B3H-4R2, Canada

**Keywords:** Choroid plexus, cuboidal epithelia, Cortical Hem, Otx1/2, Cux2, Mllt11, migration

## Abstract

The choroid plexuses (ChPs) are modified epithelial structures that penetrate all four cerebral ventricles and secrete cerebrospinal fluid. They consist of a central stroma that is vascularized with fenestrated blood vessels and connective tissue. The ChPs are of dual embryonic origin, with forebrain neuroepithelial cells contributing to the epithelial component and mesenchymal cells contributing to the stromal cells. The growth of the ChPs into the ventricular spaces is fueled by the migration and proliferation of neuroepithelial progenitor cells originating from the cortical hem. However, the genetic regulation of neuroepithelial progenitor (NEP) migration during ChP development is not well understood. Here we report the role of Mllt11 (Af1q/Tcf7c) in the formation of the ChP in part by regulating the migration of Otx1/2^+^ NEPs into the base of the fetal ChP, fueling its “treadmilling” growth into the lateral ventricle. We used *Cux2*^*iresCre/+*^ allele to selectively ablate *Mllt11* in the developing cortical hem and its principal derivatives, the ChP and hippocampus. We discovered that *Mllt11* mutants displayed thickened cuboid epithelial architecture of the ChP but maintained the epithelial organization of the outer layer of the ChP. This likely contributed to shorter lateral ventricle ChP stalks in *Mllt11* cKO brains.

## INTRODUCTION

The Choroid Plexus (ChP) of the lateral ventricle is one of the main sources of cerebrospinal fluid but its development remains a relatively understudied process. Recent work has shown that ChP formation is underlain by complex morphogenetic processes dependent on the patterning of telencephalic midline structures (Ghersi-Egea et al., 2018; Liddelow, 2015; Moore and Iulianella, 2021). The ChPs are modified epithelial structures that protrude into all four cerebral ventricles. They consist of a central stroma that is highly vascularized interdigitated with fenestrated leaky blood vessels and connective tissue (Johansson, 2014). The initial development of the ChPs begins around embryonic day (E)11 and is largely complete by E14 in the mouse, with the fourth hindbrain ventricular region ChP being the first to differentiate, followed by the lateral ventricular ChPs, and that of the remaining ventricles (Dziegielewska et al., 2001). The ChPs are of dual embryonic origin, as neuroepithelial progenitor cells (NEPs) from the cortical hem (CH) region give rise to the epithelial component and cephalic mesenchymal cells establish the stromal component (Johansson, 2014; Lun et al., 2015; Moore and Iulianella, 2021).

The differentiation of the ChP from NEPs follows a series of morphogenetic events characteristic of epithelial tissues (Moore and Iulianella, 2021). Epithelial cells change to columnar epithelium with apically located nuclei and an emerging basal connective tissue with sparse villi-like extensions. This is followed by a flattening of the outer ChP epithelial cells to become more cuboidal in shape, with centrally or apically located nuclei and more complex villi (Ek et al., 2003). Epithelial cells are sequentially added from a proliferative ventricular zone (VZ) located at the ‘root’ of each plexus as they transition through the stages outlined above (Liddelow et al., 2010). NEPs originating from the VZ are “pushed” out from the root of the ChP in the upper (dorsal) arm by newly divided cells and migrate towards the tip of the stalk. Thus, the ChP is an excellent model for epithelial differentiation during development and plays a crucial role in regulating brain function and neural stem cell maintenance (Moore and Iulianella, 2021). Yet our understanding of the genetic processes that regulate the migration of NEPs during ChP formation remains rudimentary.

Here we report a role for Mllt11/Af1q/Tcf7c (Myeloid/lymphoid or mixed-lineage leukemia; translocated to chromosome 11; also known as All1 Fused Gene from Chromosome 1q or Tcf7 co-factor) in regulating ChP formation. Work from our laboratory has identified Mllt11 as a regulator of migration of neurons in the developing cortical superficial layer, cerebellar granule cells, and retinal neuroblasts (Blommers et al., 2023; Blommers et al., 2024; Stanton-Turcotte et al., 2022; Yamada et al., 2014). During these investigations, we observed abnormal ChP morphology in the developing lateral ventricle of *Mllt11* conditional knockout (cKO) neonatal mice. Given its role in neuronal migration, we hypothesized that *Mllt11* may also control NEP migration contributing to ChP growth into the lateral ventral. We used the *Cux2*^*iresCre/+*^ driver line to inactivate *Mllt11* in the developing mouse embryonic CH, which gives rise to the lateral ventricle ChP (Fregoso et al., 2019; Moore and Iulianella, 2021; Yamada et al., 2015), using an *Mllt11*^*flox/flox*^ mouse line we previously described (Stanton-Turcotte et al., 2022). EdU birth dating and tdTomato^+^ fate mapping analysis suggests the truncated lateral ventricle ChP in the *Mllt11* null mutants was largely due to a migratory defect of Otx1/2^+^ ChP progenitors in the CH neuroepithelium. This was associated with a change in the epithelial organization of the ChP, with cells abnormally pushed inwards towards the ChP core instead of aligning into straight columns along its ventricular surface. Altogether, these findings suggest a role for Mllt11 in the migration of ChP NEPs and cytoarchitectural organization of the epithelium lining the ChP.

## MATERIALS AND METHODS

### Animals

Mice used in this study were handled in accordance with the regulations of the Dalhousie animal ethics committee and the guidelines of the Canadian Council on Animal Care. The generation of *Cux2*^*iresCre/+*^; *Mllt11*^*Flox/Flox*^: Ai9 tdTomato reporter (B6.Cg-*Gt(ROSA)26Sor*^*tm9(CAG-tdTomato)Hze*^/J, 007909, The Jackson Laboratory) conditional mouse mutants was described previously (Stanton-Turcotte et al., 2022). In the forebrain, the *Cux2* locus-driven Cre expression results in the inactivation of *Mllt11* and expression of *tdTomato* in neurogenic progenitors, superficial cortical neurons, and CH and its derivatives, including the ChP (Fregoso et al., 2019; Stanton-Turcotte et al., 2022; Yamada et al., 2015). All experiments were performed with *Cux2*^*iresCre/+*^; *Ai9; Mllt11*^*+/+*^ as the wild-type (WT) control, and *Cux2*^*iresCre/+*^; *Ai9; Mllt11*^*flox/flox*^ as the conditional knockout (cKO), harvested at E14.5 and E18.5.

### Histology and immunohistochemistry

Brains were dissected out and fixed in 4% paraformaldehyde (PFA), equilibrated in sucrose, and embedded in Optimum Cutting Temperature compound (Tissue-Tek, Torrance, CA). A total of 40 coronal sections per embryo were collected at 12µm per section. Immunohistochemistry was conducted on fetal brains as previously described (Iulianella et al., 2008; Stanton-Turcotte et al., 2022). Primary antibodies used were rabbit anti-Otx1/2 (1:500, Abcam ab21990), rabbit anti-Lhx2 (1:500; Abcam ab184337), rabbit anti-Pals1 (1:300, Proteintech 1771-1-AP), mouse anti-ZO1 (1:100, Invitrogen 339100). Donkey anti-mouse and anti-rabbit highly cross absorbed AlexaFluor 488 or 647 secondary antibodies (1:1500, Invitrogen) were used to detect primary antibodies. A cohort of sections from 4 individuals per group were stained with rat anti-Laminin (1:500, Invitrogen MA1-0600) with anti-rat 647 secondary antibodies (1:1500, Invitrogen A48265TR) and DAPI. Images were captured using a Zeiss AxioObserver inverted fluorescent microscope with x20 and x40 oil objectives and Apotome 2 Structural Illumination. Montages were assembled using Photoshop (Adobe, San Jose, CA).

### EdU (5-ethynyl-2’-deoxyuridine) labeling

10 mM EdU solution was prepared in DMSO and administered via intraperitoneal injection into dams. EdU was dosed at 30 mg/kg body in a solution of 10 mg/ml PBS. Mice were dosed at E14.5 and harvested at E18.5 and EdU staining was conducted using Click-iT EdU imaging kit (Invitrogen). The immunohistochemistry protocol was adapted such that EdU staining was performed before the addition of anti-Otx1/2 primary antibodies, as described previously (Blommers et al., 2023).

### Sampling methodology and statistics

Analysis of the CH region including the dentate neuroepithelium (DNE) and ChP stalk at E14.5 was restricted to 50 µm x 50 µm counting frames. For the ChP, the 50 μm^2^ frames were placed at the base of the ChP stalk and along the length of the stalk. Analysis of cell polarity was performed in 100µm x 50µm bins placed randomly along distal and proximal regions of the ChP. Cells positively labeled with each IHC marker were counted within counting frames or analyzed for fluorescence density within bins using ImageJ (FIJI) (Schindelin et al., 2012), and statistical analyses were performed using PRISM (GraphPad). Statistical significance was obtained by performing unpaired Student’s t-tests (two-tailed) to compare the means and standard error of the mean (SEM) between the WT and cKO data sets, with significance level set at P ≤ 0.05 (*P ≤ 0.05, **P ≤ 0.01, ***P ≤ 0.001, ****P ≤ 0.0001). For all experiments, n of 3 to 9 WT and cKO individuals were analyzed and cell counts were reported as mean ± SEM. Cell counts were performed blinded to genotype and data was presented as scatter boxplots or bar graphs with each point representing individual embryos. Analysis of Laminin thickness was performed at the distal and proximal portions of the ChP at E18.5. Three images from each individual were taken per region (I, II, and III), to capture the cytoarchitecture along the thickness of the ChP. Image J was used to acquire three thickness measurements for each image. The average thickness was calculated per image and then per individual. Statistical analysis was performed using PRISM (Graphpad). Unpaired T-tests (two-tailed) were used to compare the WT and cKO laminin thicknesses at the proximal and distal regions of the ChP. 4 WT and 4 cKO individuals were analyzed. Significance levels were P ≤ 0.05 (*P ≤ 0.05, **P ≤ 0.01, ***P ≤ 0.001, ****P ≤ 0.0001). The individual data points shown for this analysis represent the calculated average for each individual.

## RESULTS AND DISCUSSION

The CH region forms a specialized nexus at the tip of the pallial cortex that is enriched in signaling proteins and transcription factors that act to pattern telencephalon along into medial-lateral, and ventral–dorsal axes (Moore and Iulianella, 2021). *Lhx2* acts as a selector gene for cerebral cortical fate and plays a critical role in patterning the forebrain midline and CH (Bulchand et al., 2001; Chou and Tole, 2019; Monuki et al., 2001; Roy et al., 2014). We therefore used Lhx2 to assess issues with the developing dorsal telencephalic midline neuroepithelium (DNE, Fig. 1A-B). We noted an elevated number of Lhx2^+^ cells at E14.5 within the VZ in the *Mllt11* conditional knockout (cKO) relative to the wild type (WT) controls (p=0.0004; n=6; Fig. 1C). Since Lhx2 is critical in establishing an accurate boundary region between the dorsal telencephalon and CH, we examined whether there were any notable changes to the CH region by staining for the homeodomain transcription factor Otx2, which regulates the formation the anterior-most forebrain regions, including the CH (Johansson et al., 2013). As *Otx2* expression is mainly in the diencephalon, mesencephalon, choroid plexus, and the hippocampal anlage while *Otx1* expression is highest in the forebrain proliferative layers of the neocortex (Acampora et al., 2000; Larsen et al., 2010), we used an antibody that detects both Otx1 and Otx2 (Otx1/2) to demarcate NEPs in the CH and ChP. We noted that the CH area, outlined by the hatched yellow line (Fig. 1D-F), was significantly smaller in the *Mllt11* cKOs relative to control fetal brains (p=0.003; n=6). This suggested that *Mllt11* loss led to a slightly reduced CH region, which may have affected the numbers of NEPs that migrate during ChP growth. The conditional strategy removed *Mllt11* and activated *tdTomato* expression from the Ai9 reporter mouse allele (Stanton-Turcotte et al., 2022), allowing for the fate-mapping of Cux2^+^ cells in the CH and its derivatives, including the entire ChP stalk (Fig.1G, H) (Fregoso et al., 2019; Stanton-Turcotte et al., 2022; Yamada et al., 2015). We observed decreased tdTomato^+^ cell staining along the base of the developing ChP stalk in *Mllt11* cKO mutants relative to WTs (white arrowheads, Fig. 1G, H), suggesting a NEP migration defect.

**Figure 1.**
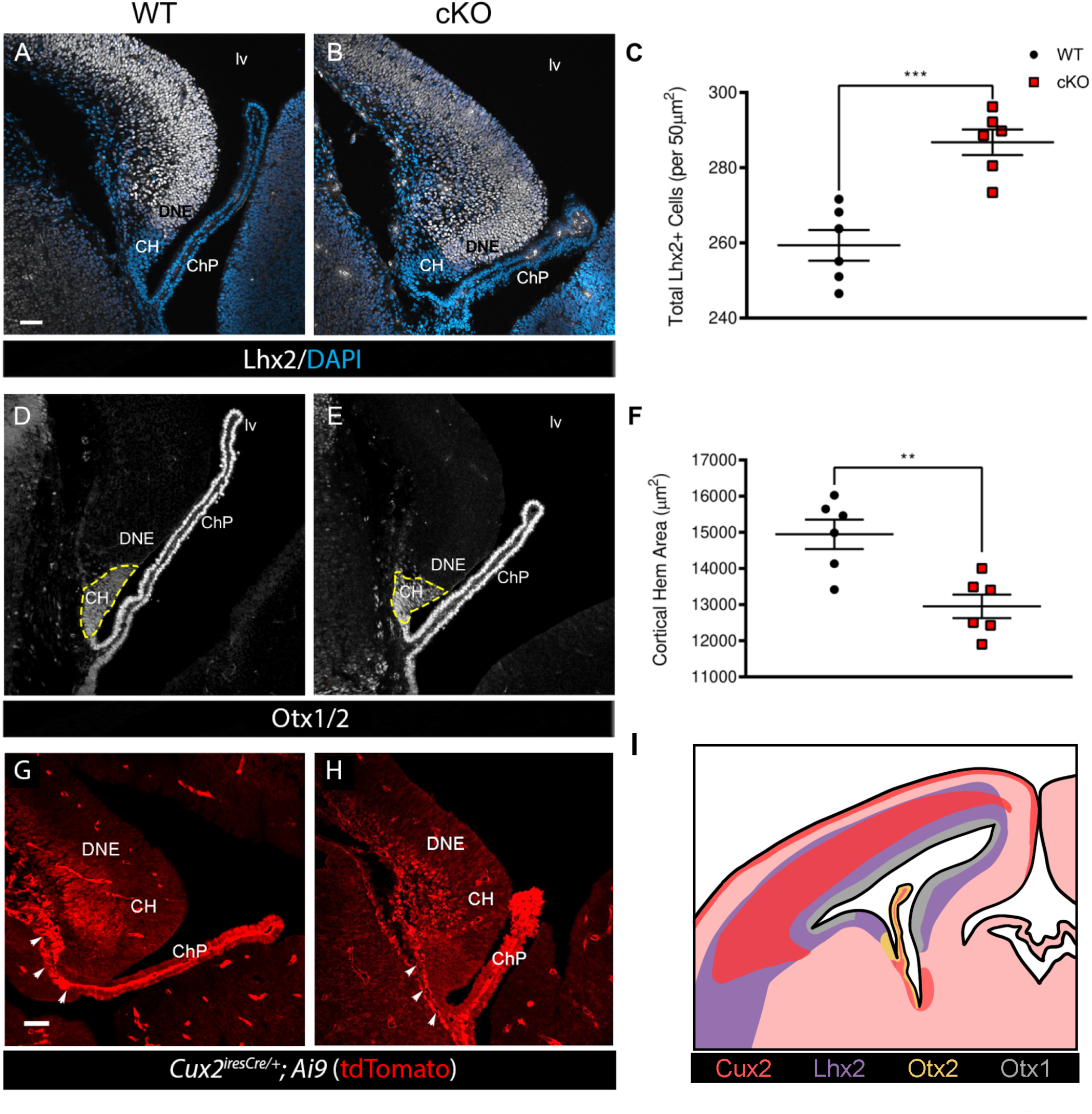
*Mllt11* loss altered development of dorsal telencephalic midline structures. (A, B) Coronal sections of DTM region at E14.5 WT control vs. *Cux2*^*iresCre/+*^driven *Mllt11* cKO stained for Lhx2 (white) counterstained with DAPI (blue). (C) Scatter boxplot identifying the total number of Lhx2^+^ cells per 50 μm^2^. *Mllt11* cKOs displayed a significant increase in Lhx2^+^ cells in the dorsal medial telencephalon relative to WTs (p=0.0004, n=6). (D, E) Coronal sections of medial telencephalon of E14.5 WT vs. cKO stained for Otx1/2 (white) to label the CH region and ChP. (F) Scatter boxplot identifying the total CH area in μm^2^. A significant decrease is observed in the CH area in the cKO in comparison to WT controls (p= 0.003; n=6). (G, H) Coronal sections of forebrains from E14.5 WT vs. cKO embryos with tdTomato (red ; *R26r*^*tdTpmato/tdTomato*^ Ai9 reporter) fluorescence as a readout for *Cux2*^*iresCre/+*^-mediated recombination in the CH and ChP stalk. White arrowheads depict tdTomato^+^ fate-mapped NEPs contributing to ChP formation. (I) Representative diagram of *Cux2*^*iresCre/+*^-driven fate mapping, and Lhx2, Otx1 and Otx2 expression in the fetal mouse brain. Quantification of data presented as mean ± SEM. Abbreviations: CH, cortical hem; ChP, choroid plexus; DNE; dentate neuroepithelium; hf, hippocampal fissure, lv, lateral ventricle.

### *Mllt11* Loss Decreased ChP Stalk Length

To evaluate the effect of *Mllt11* loss on ChP development, we focused our analysis at E14.5 when ChP formation in the lateral ventricle peaks and used Otx1/2 immunostaining to outline the CH and ChP in our morphometric analyses (Johansson et al., 2013). For each individual, three images of the ChP were acquired (Fig. 2). The ChP stalk length was significantly shorter in the *Mllt11* cKOs (407.1 ± 22.95μm; n=9) relative to WTs (540.1 ± 27.04; p<0.0001; Figure 2A-C). *Mllt11* cKO mutants displayed thicker ChP stalks in fate-mapping and immunostaining experiments (Figs. 1H, 2B), and we noted an average increase of ChP stalk thickness of approximately 3μm, which did not reach statistically significant difference between cKOs (29.33 ± 1.57μm; n=9) and WT controls (26.17 ± 0.93μm; n=9; p=0.10; Fig. 2A, B, D). This suggested that *Mllt11* loss from CH NEPs led to altered ChP neuroepithelial cytoarchitecture, but the cellular mechanisms for this phenotype was unclear. We observed an abnormal clustering of Otx1/2^+^ cells at the base of ChP along the ventricular surface relative to the distal tip 350 µm away (Fig. 2E), likely reflecting aberrant migration of *Mllt11* mutant NEPs (Fig. 2E).

**Figure 2.**
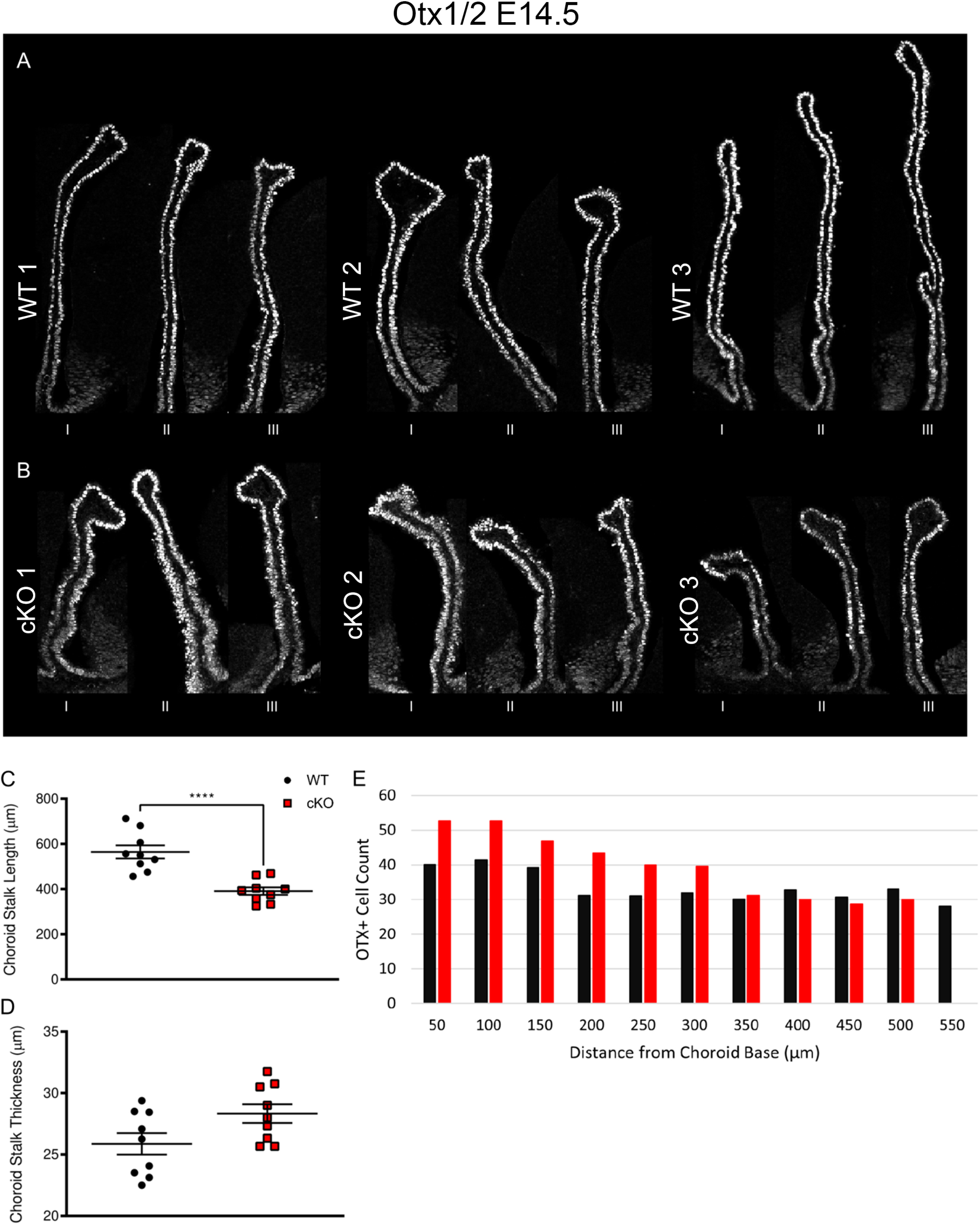
*Mllt11* loss resulted in truncated telencephalic ChP. (A, B) Cross sections of telencephalic ChPs from 3 different individual WT controls and *Mllt11* cKO fetuses at E14.5 stained for Otx1/2 to label the ChP. Three images were acquired per individual, labeled as I, II, III, reflecting the thickness of the ChP. (C) Scatter boxplot of ChP stalk length in μm. ChP stalk length was significantly shorter in the cKOs (407.0 ± 22.95μm; n=9) relative to WTs (540.1 ± 27.04μm; n=9; P=0.0018). (D) Scatter boxplot quantifying ChP stalk thickness in μm. cKO fetuses (29.33 ± 1.57μm, n=9) displayed no significant difference in ChP thickness relative to WTs (26.17 ± 0.93μm; n=9; P= 0.10). (E) Bar chart quantifying total Otx1/2^+^ cells per 50 μm^2^, beginning at the base of the ChP stalk and counted in 50μm increments. Quantification of data presented as mean ± SEM. Abbreviations: ChP, Choroid Plexus, SEM, standard error of the mean.

### *Mllt11* loss altered epithelial cell contacts along the ChP stalk

The aberrant cellular morphology of the ChP stalk in *Mllt11* cKOs fetal brains led us to examine the epithelial organization of the stalk using *Zona Occludens-*1 (ZO-1) antibodies, which labels the basement membrane. ZO-1 is normally expressed in epithelial and endothelial cells and acts as a scaffold protein cross-linking and anchoring tight junctions (McNeil et al., 2006). Importantly, we noted a marked change in the typical cuboidal epithelial organization of the maturing ChP stalk in the E18.5 *Mllt11* cKOs, with mutant cells clustering aberrantly along the width of the ChP (Fig. 3C; arrowheads in Fig. 3D), instead of being arrayed in single file along the ChP stalk length as in WT controls (Fig. 3A, B). The *Mllt11* mutant ChP displayed more bridging between displaced individual cells, outlined by the ectopic ZO-1 boundary staining along the width of the ChP (arrowheads, Fig. 3D). This suggested that *Mllt11* loss affected the formation of epithelial cell boundaries of the ChP stalk.

**Figure 3.**
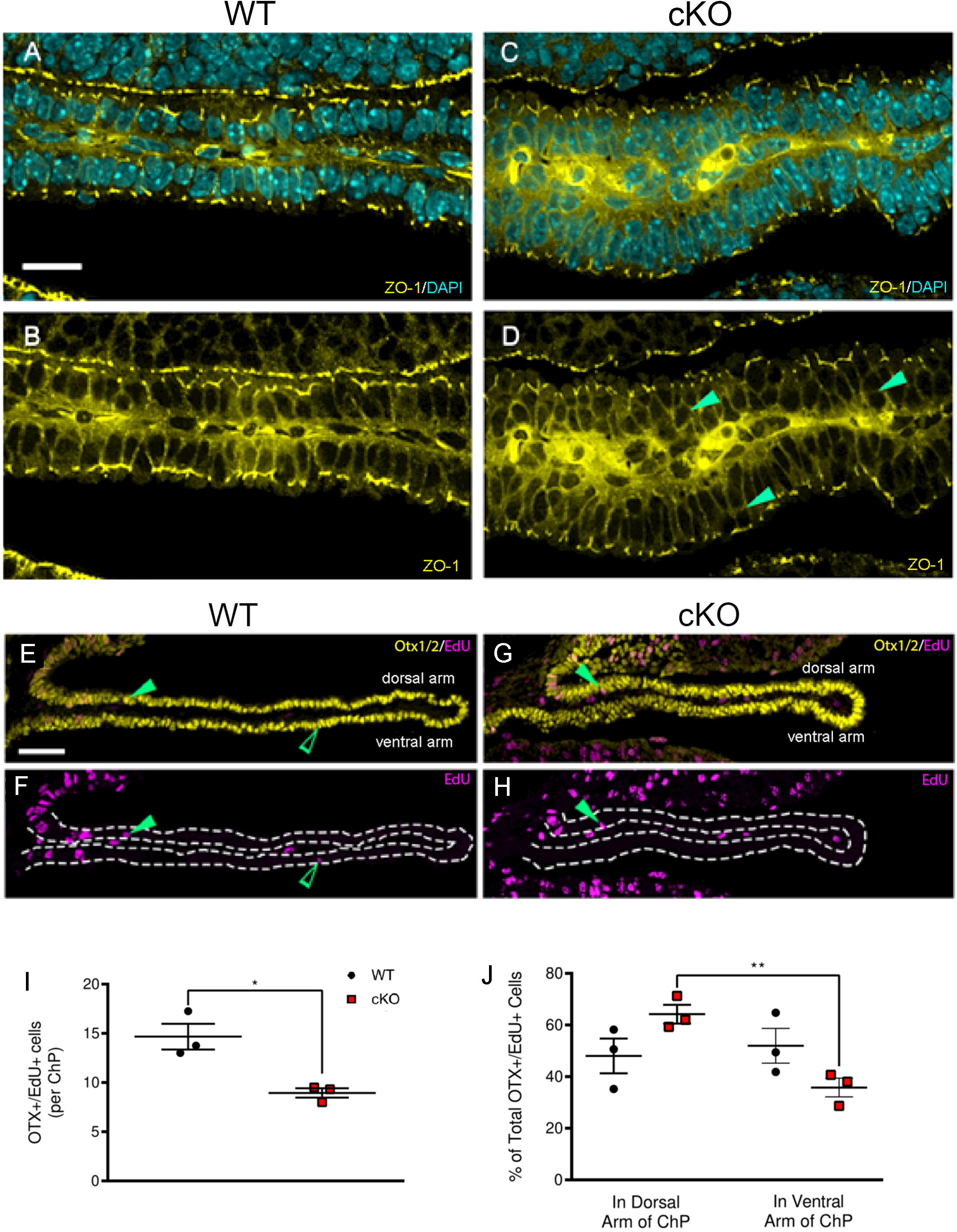
Reduced treadmilling migration of Otx1/2^+^ ChP progenitors in *Mllt11* cKOs. (A-D) Cross-sectional view of WT and *Mllt11* cKO telencephalic ChP stained for ZO-1 (yellow) and DAPI (blue). Solid green arrowheads (D) identify the disorganized placement of cKO ChP epithelial cells relative to the single cell layer organization of WT ChP. (E-H) Cross section of WT and cKO ChP strained for Otx1/2 (yellow) and EdU (violet) at E18.5 (EdU pulse at E14.5). Solid and open green arrowheads identify Otx1/2^+^/EdU^+^ cells in the dorsal arm and the ventral arm of the stalk, respectively. (I) Scatter boxplot identifying Otx1/2^+^/EdU^+^ cells per ChP stalk, reporting a significant decrease in cKOs (p= 0.015, n=3) following an EdU pulse at E14.5 and analysis at E18.5. (J) Scatter boxplot identifying the percent total of Otx1/2^+^/EdU^+^ cells in the dorsal arm of the stalk and the ventral arm of the stalk, reporting a significant decrease in percent of Otx1/2^+^/EdU^+^ in the ventral arm of the cKOs relative to the dorsal arm (p= 0.005, n=3) at E18.5. Scale bar = 50μm. Quantification of data presented as mean ± SEM. Abbreviations: ChP, Choroid Plexus.

### Aberrant migration of ChP NEPs in *Mllt11* cKO mutants

ChP growth is fueled by a ‘treadmilling” behavior whereby migratory ChP stalk progenitor cells generated from NEPs invade the developing ChP base and move its dorsal to ventral axis (Moore and Iulianella, 2021). To directly assess whether the truncated ChP in the *Mllt11* mutants reflected reduced migration of Otx1/2^+^ NEPs from the CH ventricular region, we pulsed dams with a single pulse of the DNA analog EdU at E14.5 to label newly divided NEPs contributing the ChP formation and assessed birth dating at E18.5. We observed a significant decrease in the overall numbers of Otx1/2^+^/EdU^+^ cells in the cKO ChP at E18.5 (p=0.015, n=3; Fig. 3E-J). Moreover, cKOs displayed decreased EdU^+^ NEPs populating the distal tip of the ChP stalk (green open arrow, Fig. 3F). The ChP is composed of two neuroepithelial arms: the dorsal and ventral. NEPs migrate from the CH ventricular region and first enter the dorsal arm of the ChP (solid green arrowheads, Fig. 3F), to eventually loop around and invade the ventral arm (open green arrowheads, Fig. 3F). If *Mllt11* loss affects NEP migration, then we expect to find reduced numbers of EdU^+^ entering the ventral arm of the ChP at fetal stages. We quantified the numbers of Otx1/2^+^/EdU^+^ in the dorsal and ventral arms and found greater numbers of double-labeled cells in the dorsal arm of the *Mllt11* cKO ChP, along with a corresponding decrease in the ventral arm (Fig. 3J). This demonstrated that *Mllt11* loss led to reduced treadmill migration of NEPs along the ChP stalk.

### *Mllt11* cKO mutants did not exhibit altered epithelial polarity within the lateral ventral choroid plexus

*Mllt11* loss led to a migration defect in ChP NEPs contributing to a shortening of the ChP stalk length. ZO-1 staining revealed that the *Mllt11* cKO mutants also displayed disorganized epithelial organization in the ChP stalk (arrowheads, Fig. 3D), with cells displaced centrally towards the core of the ChP stack instead of organized in a single layer of cuboidal epithelium (Fig. 3C), creating a juxtapositioning of cells along the ChP stalk length. This may reflect a migration defect or alternatively be due to a loss of epithelial polarity and cuboidal cytoarchitecture. To explore this possibility we stained for Pals1 (Protein associated with Lin Seven 1), a protein that is asymmetrically expressed in the outer membrane of epithelial cells, including the ChP (Christensen et al., 2018; Ozcelik et al., 2010). The overall organization of Pals1 staining in the lateral ventricle ChP epithelium was similar in both control and *Mllt11* cKO, with staining restricted to the outer membrane of the epithelial cell layer along the ventricular edge of the choroid stalk (Fig. 4A). This indicated the epithelial polarity was maintained in the cKOs. We confirmed an abnormal thickening of the outer choroid epithelium in the *Mllt11* cKOs, with epithelial cells aberrantly displaced towards the vascular core (insets, lower panel, Fig. 4A). There was a trend towards reduced proximal ChP area staining for Pals1 in the cKOs, although the differences were not statistically significant (Fig. 4B, C). Our morphometric analyses confirmed significantly increased thickness in the proximal ChP stalk of *Mllt11* cKO mutants (24.99μm ± 1.37; n=4, vs. controls: 15.08μm ± 0.81; n=4; p= 0.002; Fig. 4F), but not distal stalk (Fig. 4G), and explained why the average ChP thickness did not reach statistical significance (Fig. 2D). It is likely that difference in thickness along the proximo-distal axis of the stalk is at least in part due to altered NEP migration entering the base of the ChP. This aberrant accumulation of cells in the proximal stalk was supported by reduced tdTomato^+^ fate labeling in the Mllt11 cKOs (Fig. 1G, H) and was accompanied by reduced Otx1/2^+^/EdU^+^ double-positive cells progressing from the dorsal to ventral halves of the neonatal ChP (Fig. 3E-J). Our findings indicate that the *Mllt11* mutant choroid phenotype is primarily restricted to the epithelial cell layer, impacting both the migration of NEPs fueling the growth of the ChP stalk, as well as possibly affecting the transition of the epithelial cells from columnar to cuboidal organization, leading a thickened and disorganized choroid epithelia cell layer.

**Figure 4.**
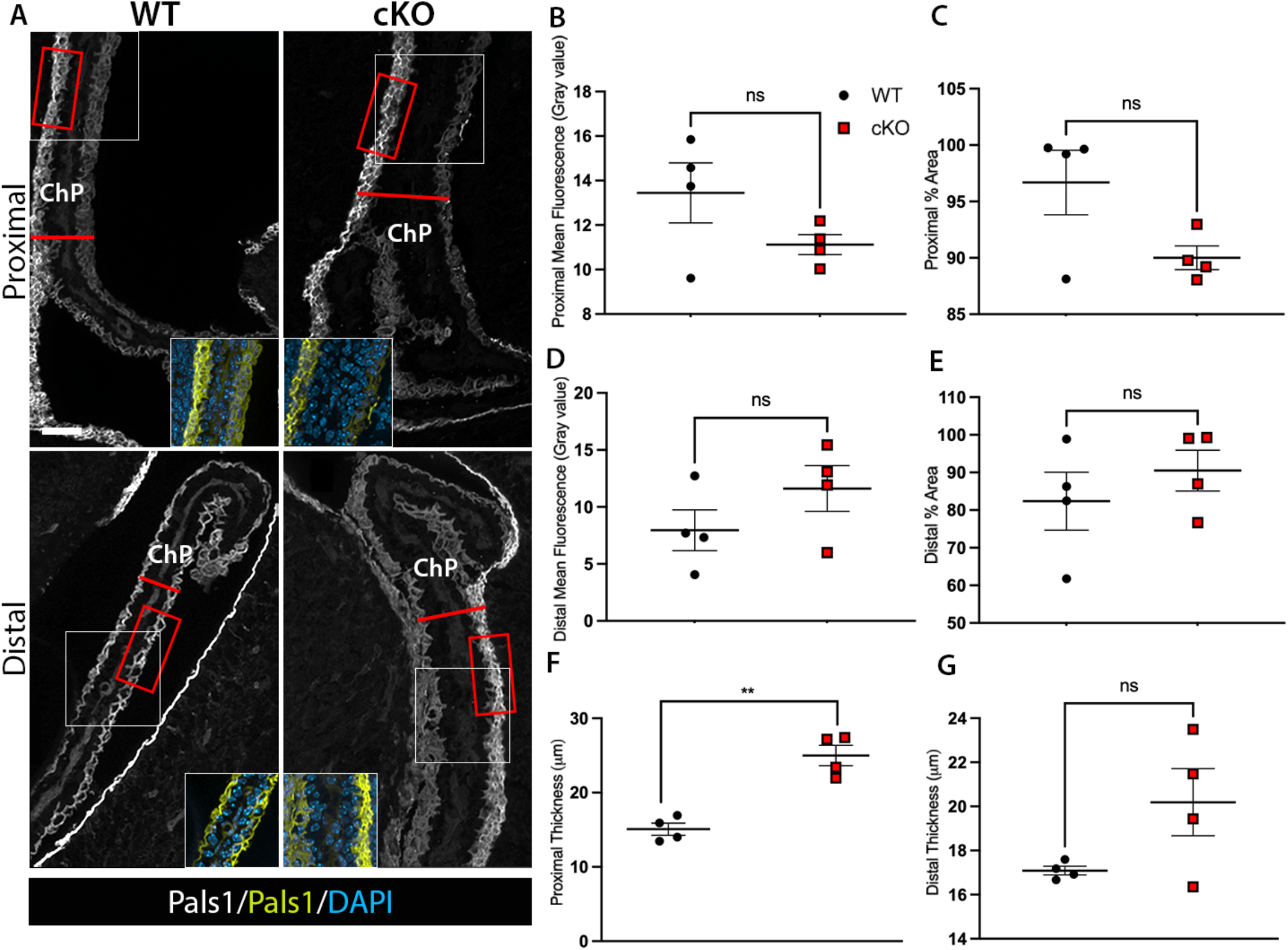
*Mllt11* cKOs displayed a thickened the epithelial layer along the ChP stalk. Coronal sections of the proximal and distal regions of the ChP in WT and *Mllt11* cKO mice stained for Pals1. (B-E) Bar graphs depicting no significant differences in mean fluorescence levels of or percentage of area containing Pals1 staining respectively, quantified proximally (B, C) or distally (D, E). (F, G) Scatter boxplots quantifying a significant increase in ChP thickness (F) proximally in *Mllt11* cKO (24.99μm ± 1.37; n=4) compared to WT (15.08μm ± 0.81; n=4; P=.002), with no significant differences seen (G) distally. White boxes indicate sites of insets. Red boxes indicate example regions sampled for analysis of immunofluorescence. Red lines indicate sample regions measured for thickness analysis. Scale bar = 50μm. Quantification of data presented as mean ± SEM. Abbreviations: ChP, choroid plexus.

To determine if the thickened epithelial cell layer in *Mllt11* cKOs was due to changes in epithelial cytoarchitectural organization, we next explored the integrity of the basal lamina, which anchors an connects the ChP epithelial cells (Gabrion et al., 1998; Peraldi-Roux et al., 1990, Vandenbroucke et al., 2012). We investigated this by staining for the basal lamina of the ChP epithelium in E18.5 *Mllt11* cKOs vs WT controls using Laminin antibody staining (Fig. 5). Images were acquired in both the proximal (n=4 per genotype; Fig. 5A-B) and distal regions (Fig. 5C-D) of the ChP. We saw no differences in the width of the Laminin staining at the proximal or distal regions of the ChP (Fig. 5E-F; p > 0.05). This demonstrated that the *Mllt11* mutant ChP maintained its overall epithelial morphology and tissue relationship with the vascular core. The aberrant juxtapositioning of epithelial cells revealed by ZO-1 and Pals1 staining may instead reflect abnormal NEP migration and/or altered maturation of epithelial cells.

**Figure 5.**
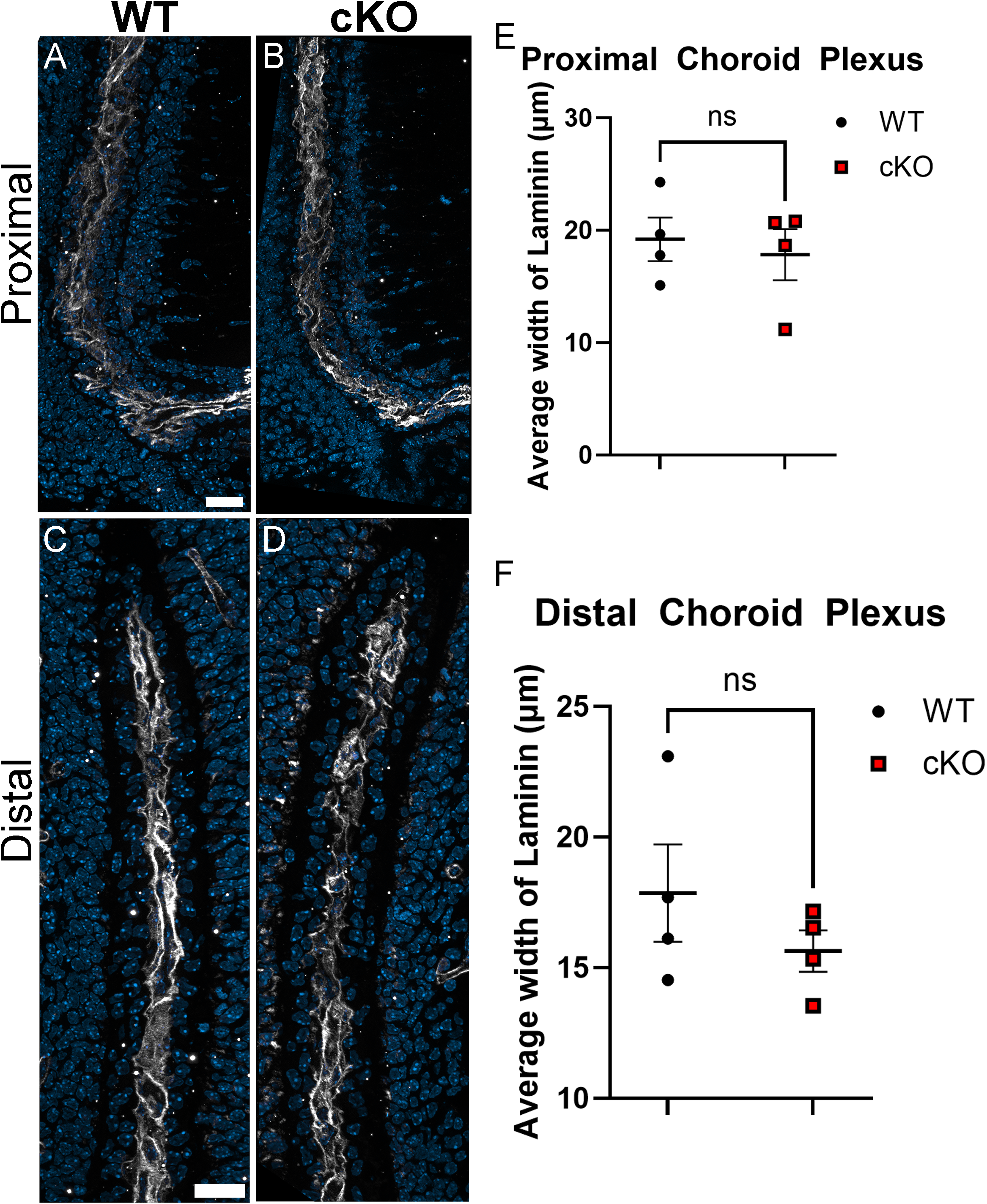
Cuboidal epithelial cytoarchitecture was intact in *Mllt11* cKOs. (A-D) Representative images of coronal sections of both the proximal and distal regions of the ChP in E18.5 WT and *Mllt11* cKO mice were stained for Laminin to label the basal lamina of the epithelial cell layer. (E) Scatter box plot quantifying the average width of the laminin stain of the proximal region of the choroid plexus. (F) Scatter box plot quantifying the average width of the laminin stain of the distal region of the choroid plexus. Quantification of data presented as mean ± SEM. Scale bar in A, C = 20μm.

## CONCLUSION

The ChP epithelium develops from CH region generating a highly organized tissue that acts as an interface between the peripheral circulation and the CNS (Ek et al., 2003; Grove et al., 1998; Liddelow et al., 2010). We now report that *Mllt11* loss from the CH neural progenitor region resulted in a significantly truncated ChP with notable defects in its epithelial morphology. In mouse fetuses the ChP epithelium consists of structures that vary from simple columnar to cuboidal organization (Huang et al., 2009; Liddelow et al., 2010). The ChP undergoes an intriguing distal-proximal maturation process whereby proliferative cells are added to the root of the plexus, forming in a “conveyor belt”-like or “treadmilling” manner (Liddelow et al., 2010). A small population of proliferative cells within the root continue to divide and contribute to the epithelial component of the ChP along the length of the stalk (Huang et al., 2009; Liddelow et al., 2010). As the epithelium progresses from the proliferative progenitor population along the ventricular surface of the CH region, it transitions from a columnar epithelium into a highly polarized cuboidal epithelium necessary for important barrier functions (Ek et al., 2003; Liddelow et al., 2010). However, *Mllt11* cKOs displayed a slightly thickened layer epithelial cells along the length of the ChP stalk, and shortened ChP length. The mutant epithelial cells were abnormally displaced along the short axis of the ChP, creating ectopic cell boundaries in the epithelial layer, which were outlined by the basement membrane marker ZO-1 and the asymmetric determinant Pals1. Despite the abnormal appearance of the *Mllt11* mutant ChP, the epithelial layer maintained contact with the vascular core, as revealed by the largely normal Laminin staining outlining the basal lamina of the ChP epithelium.

Additionally, EdU birth dating experiments revealed reduced numbers of cells reaching the ventral arm of the ChP stalk in the cKOs, suggesting a defect in the migration of newly born Otx1/2^+^ NEPs, which ultimately translated into reduced stalk length. The reduced ChP length in *Mllt11* mutant brains could also be due to the reduced Otx1/2^+^ CH progenitor region, which would lead to fewer NEPs invading the growing ChP. This phenotype was consistent with the reduced tomato^+^ NEP region in *Mllt11* cKOs, as revealed by the *Cux2*^*ireCre/+*^; *Ai9* tdTomato^+^-fate mapping data. Interestingly, the reduced CH region was associated with an expanded Lhx2^+^ DNE region in the cKOs, suggesting that Mllt11 may regulate CH vs DNE fate choice; the molecular mechanism of this regulation is currently unknown. Lastly, the shortened ChPs may also be due to the altered epithelial cytoarchitecture of cKO stalks, with a thickened epithelial cell layer reflecting aberrantly juxtaposed towards the core of the ChP stalk, as revealed by the ZO-1 and Pals1 staining. The altered migratory behavior of *Mllt11* mutant Otx1/2^+^ CH NEP progenitors is consistent with the emerging view that Mllt11 is a novel cytoskeletal-associated protein required for neuronal migration in the cortex (Stanton-Turcotte et al., 2022), retina (Blommers et al., 2023), and cerebellum (Blommers et al., 2024).

## Abbreviations

(CH): Cortical Hem
(ChP): Choroid Plexus
(SVZ): Subventricular Zone
(NEPs): Neuroepithelial Progenitors.

## ACKNOWLEDGMENTS

We gratefully acknowledge funding from the National Research and Engineering Council of Canada (NSERC RGPIN-2020-03925). SM was supported by a CIHR MSc award.

